# Theta oscillations tag memories for sleep-dependent consolidation

**DOI:** 10.1101/2025.01.25.634851

**Authors:** Dan Denis, Zhiyi Chen, Manroop Kaur, Ben Clayden, Thomas Schreiner, Scott A. Cairney

## Abstract

How does the brain select which experiences to consolidate into long-term memory? Numerous neurobiological frameworks suggest that certain memories are “tagged” at learning for consolidation during later sleep. However, experimental evidence of such a tagging mechanism in the human brain is lacking. Using electroencephalography in humans, we reliably differentiate brain states for memories that are tagged at learning for consolidation across sleep or wakefulness. The tagging of memories for consolidation across sleep (but not wakefulness) was linked to 3-8Hz theta rhythms during learning. The magnitude of this tagging-related theta response predicted the coupling of slow oscillations to sleep spindles during post-learning sleep (an established neural correlate of sleep-dependent memory processing). In turn, slow oscillation-spindle coupling was associated with better memory performance at the post-sleep test. These findings provide new insights into the neural mechanisms through which our brains determine which information is retained for the future.

## Introduction

Our memories are not an unedited replay of daily events, but rather a showreel of our most salient experiences. This adaptive filtering of memory is thought to be driven by a “tagging” process at learning, such that certain memories are prioritised for consolidation during offline periods [1,2]. Selective memory processing is considered essential to prevent system overload and ensure that memories are preserved in accordance with our cognitive and emotional goals.

Sleep has long been implicated in memory consolidation and may play a unique role in the selective retention of memories tagged at prior learning [3]. Indeed, there is evidence that salience cues at encoding prompt selective memory processing during sleep; information that is perceived as emotionally arousing [4–6], linked to financial reward [7,8] or considered relevant for future events [9–11] appears to be preferentially strengthened in the sleeping but not waking brain (though see [12]). Numerous theoretical frameworks therefore position sleep as a point of memory triage, with memories being consolidated on the basis of neurobiological tags established at the initial learning phase [1–3,13,14].

Findings from recent animal studies are consistent with the idea that memories are tagged at learning for overnight consolidation. Huelin Gorriz et al (2023) found that sleep replay of hippocampal place cells increased with the frequency of maze traversals during prior wakefulness, but decreased when the experience was already familiar, suggesting that repetition and novelty influence which memories are tagged for sleep-dependent memory processing [15]. Relatedly, Yang et al (2024) found that sleep sharp-wave ripples (SWRs) replayed maze trajectories that were reactivated during wake SWRs linked to reward, characterising a mechanism for selecting which memories are consolidated in later sleep [16]. However, evidence for an analogous mechanism in humans has yet to be established and represents an important translational gap in our understanding of sleep’s role in memory.

If the human brain tags certain memories for selective strengthening during later sleep, then it should be possible to differentiate between brain activity patterns at learning for memories that are specifically tagged for consolidation over sleep, relative to those that are not. Multivariate neuroimaging analyses can meet this need by decoding brain states that are based on distinct patterns of neural activity [17]. In the current context, memories retained across sleep-filled delays should be distinguishable from memories retained across wake-filled delays, based only on the neural operations (i.e., tags generated) at initial learning. We tested this hypothesis in the present study by employing multivariate classifiers to differentiate between encoding trials for memories that were remembered after sleep and encoding trials for memories that were remembered after an equivalent period of wakefulness.

Although multivariate analyses can provide evidence of mnemonic tagging, they do not address the specific oscillatory rhythms that underpin this process. Theta oscillations (∼3-8Hz) are a promising candidate brain rhythm that may set the scene at learning for selective memory processing during later sleep. In humans, larger overnight memory gains are predicted by higher levels of theta activity during pre-sleep learning [18], mirroring animal research linking theta activity at wakefulness to replay during sleep [15]. Complementing these findings, other work has shown that the presence of reward or arousal cues at learning modulates theta power in a manner that is predictive of subsequent memory [19,20]. Here, we isolated frequency specific activity in the theta band at learning to test the hypothesis that theta oscillations uniquely support the tagging of memories for sleep-dependent consolidation.

If theta oscillations at learning act as an instructional cue for overnight memory processing, they should also influence the neural signatures of sleep-dependent memory consolidation. Memory representations are thought to be reactivated during sleep through tightly coupled interactions between global slow oscillations (SOs; ∼1Hz), thalamocortical sleep spindles (∼12-15Hz) and hippocampal ripples [21–25]. Growing evidence indicates that these sleeping brain rhythms, particularly sleep spindles, promote the selective strengthening of future relevant information [8,9,26], and are correlated with theta power at learning [18]. Building on this work, we examined whether the magnitude of tagging-related activity (i.e., theta rhythms) at learning predicts the emergence of SO-spindle coupling during later sleep, with increases in SO-spindle coupling subsequently predicting sleep-related gains in performance.

In the current study, we set out to test for the existence of a tagging mechanism for sleep-dependent memory consolidation in humans. Participants completed a two-visit, within-subjects experiment where they learned word-object pairings and were tested before and after a 2-h sleep opportunity (nap) or an equivalent period of wakefulness. Using multivariate analysis of electroencephalography (EEG) data, we found evidence that brain activity patterns at learning differentiate between memories remembered after sleep from memories remembered after wake. Moreover, theta power at learning was increased for memories retained across sleep but not wakefulness, with the magnitude of this theta response predicting the emergence of SO-spindle coupling during later sleep, which in turn predicted memory retention. Together, these results are consistent with a dynamic tagging of memory for sleep-dependent consolidation that is curated by theta oscillations at learning.

## Results

Thirty-one healthy young adults (M_age_ = 20 years, range = 18-23, 68% female; **Table S1**) took part in the study. Participants began each visit by learning a set of word-object pairs (**Figure 1A**). Immediately after learning, and again following a 2-h delay (**Figure 1B**), participants were presented with the words in isolation, one after another, and for each word were instructed to recall the associated object. To minimise any effects of retrieval practice on our behavioural measures of memory consolidation [27,28], participants were tested on different subsets of word-object pairs at the immediate and delayed tests. The 2-h delay was filled with either a daytime nap (**Table S2** for sleep architecture) or time spent awake in the sleep laboratory watching nature documentaries (delay condition order counterbalanced).

**Figure 1.**
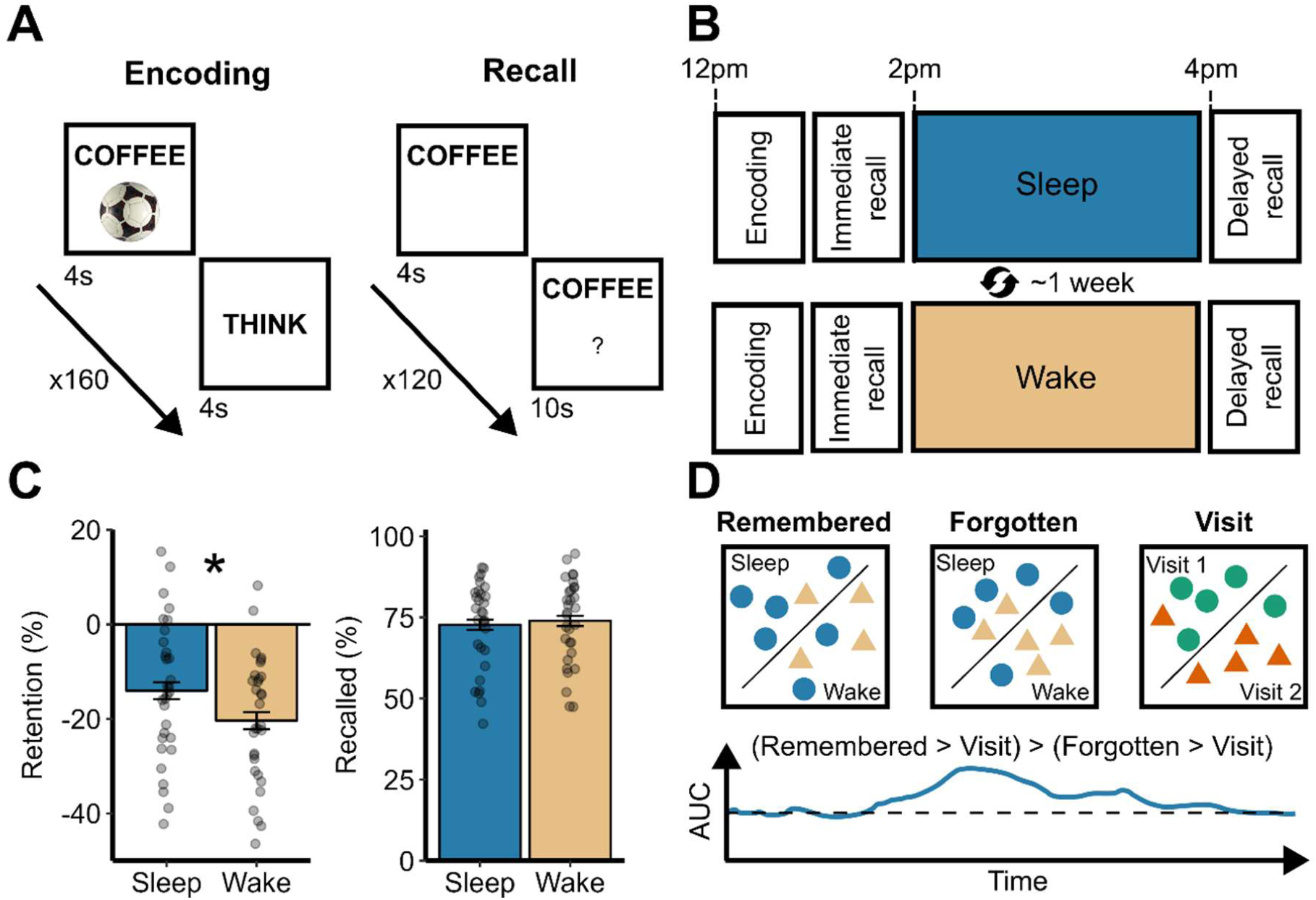
Experimental design. **A** – During learning, participants were presented with 160 word-object pairs. At recall, memory was probed by way of a cued recall test. **B** – Memory was tested immediately (immediate recall) and again following a 2-h delay of either sleep or wake (delayed recall). Sleep and wake visits were manipulated within-subjects and performed one week apart in a counterbalanced order. **C** – Retention (the change in performance at delayed compared to immediate recall, with a more negative number indicating more forgetting) was significantly better over the sleep delay compared to the wake delay (* = *p* < .05). There was no sleep/wake visit difference in proportion of items correctly recalled at immediate recall. Error bars indicate the standard error. **D –** Multivariate classification approach. Multivariate classifiers were trained on the EEG time series at learning to distinguish trials that were later remembered after sleep from trials that were later remembered after wake. As control analyses, additional classifiers were trained to distinguish between trials that were forgotten in the sleep and wake conditions, and also to distinguish between experimental visits 1 and 2 (independent of the sleep or wake condition). For statistical analysis, a double subtraction was performed, meaning any classifier values significantly above zero reflected a memory-specific signature distinguishing between trials remembered after sleep and trials remembered after wake, independent of any non-specific session differences (AUC: area under the curve, the time series shown here is an illustrative example).

Confirming the presence of a behavioural effect of sleep for memory, retention was better when participants slept relative to when they remained awake (t (30) = 2.49, p = .019, d = 0.45; **Figure 1C, left**). Memory performance did not differ between immediate tests that took place before the sleep or wake delays (t (30) = -0.53, p = .60, d = -0.10; **Figure 1C, right**). There was no effect of visit number (visit 1 vs 2) on either immediate memory performance (t (30) = -2, p = .055, d = -0.36) or retention (t (30) = 0.16, p =.87). Memory scores are shown in **Table 1**. Given the observed sleep benefit, we next turned our attention to tagging mechanisms at learning that pre-select memories for sleep-dependent consolidation.

**Table 1.** Memory scores.

Memory scores
|  | Immediate recall |  | Delayed recall |  | Retention |  |
| --- | --- | --- | --- | --- | --- | --- |
|  | M | SD | M | SD | M | SD |
| <u>Hit rate</u> |  |  |  |  |  |  |
| Sleep | 90.92 | 5.96 | 85.81 | 7.09 |  |  |
| Wake | 89.93 | 5.97 | 80.39 | 7.09 |  |  |
| <u>False alarm rate</u> |  |  |  |  |  |  |
| Sleep | 5.21 | 5.43 | 6.08 | 4.83 |  |  |
| Wake | 5.15 | 5.52 | 6.12 | 4.83 |  |  |
| <u>Associative memory</u> |  |  |  |  |  |  |
| Sleep | 72.74 | 8.65 | 62.84 | 8.87 | -12.33 | 8.96 |
| Wake | 73.91 | 8.65 | 59.15 | 8.88 | -19.88 | 8.86 |

*Note*, M = mean, SD = standard deviation. Associative memory calculated as % of correctly identified old words for which the associated image was recalled. Retention calculated within-subject as [(delayed recall – immediate recall) / immediate recall].

### Brain activity patterns at learning differentiate memories that are retained over sleep or wake

If memories are selectively tagged at learning for consolidation during sleep, then it should be possible to distinguish between memories that are remembered after sleep or wakefulness based on unique patterns of brain activity at learning. To this end, we applied a multivariate classifier to the EEG data acquired at learning to distinguish between trials that were later recalled after sleep from trials that were later recalled after wake. To isolate signatures of mnemonic tagging for subsequent sleep vs wake-based memory processing, whilst accounting for non-specific between-session differences at learning, three separate classifiers were used. A *Remember* classifier was trained to distinguish between trials remembered following sleep compared to trials remembered following wake. A *Forgotten* classifier was trained to distinguish between trials forgotten after sleep compared to trials forgotten after wake. Finally, a *Visit* classifier was trained to distinguish learning at visit 1 from learning at visit 2 (independent of when the sleep or wake conditions occurred). Compared to surrogate decoders trained on randomised condition labels [25], all three classifiers achieved significant above chance classification (all *p_cluster_* < .003, see **Figure S1** for classification time series).

We then isolated unique brain activity patterns at learning for memories that were subsequently recalled after sleep or wakefulness via a double-subtraction approach. First, the time series for the *Visit* classifier was subtracted from the time series for both the *Remembered* and *Forgotten* classifiers, generating visit-corrected signals of successful and unsuccessful learning, respectively. The corrected *Forgotten* classifier was then subtracted from the corrected *Remember* classifier, with the resultant signal distinguishing memories retained over sleep from memories retained over wakefulness (see the illustrative example in **Figure 1D**).

Significant classification was achieved 0.87-1.13s post stimulus onset (t_sum_ = 129, p_cluster_ = .014, d_cluster_ = 0.72; **Figure 2A**). This shows that brain activity patterns linked to successful learning differed according to whether the memory was later remembered across sleep or wake. Classifiers trained on trials that were subject to testing immediately after learning (i.e., *before* the sleep or wake delay) could not reliably differentiate between the sleep and wake conditions (t_sum_ = 78.36, p_cluster_ = .10; **Figure S2**). Taken together, these findings are consistent with the idea that memories are tagged at learning for selective consolidation during later sleep. We next turned our attention to the brain rhythms supporting this tagging process.

**Figure 2.**
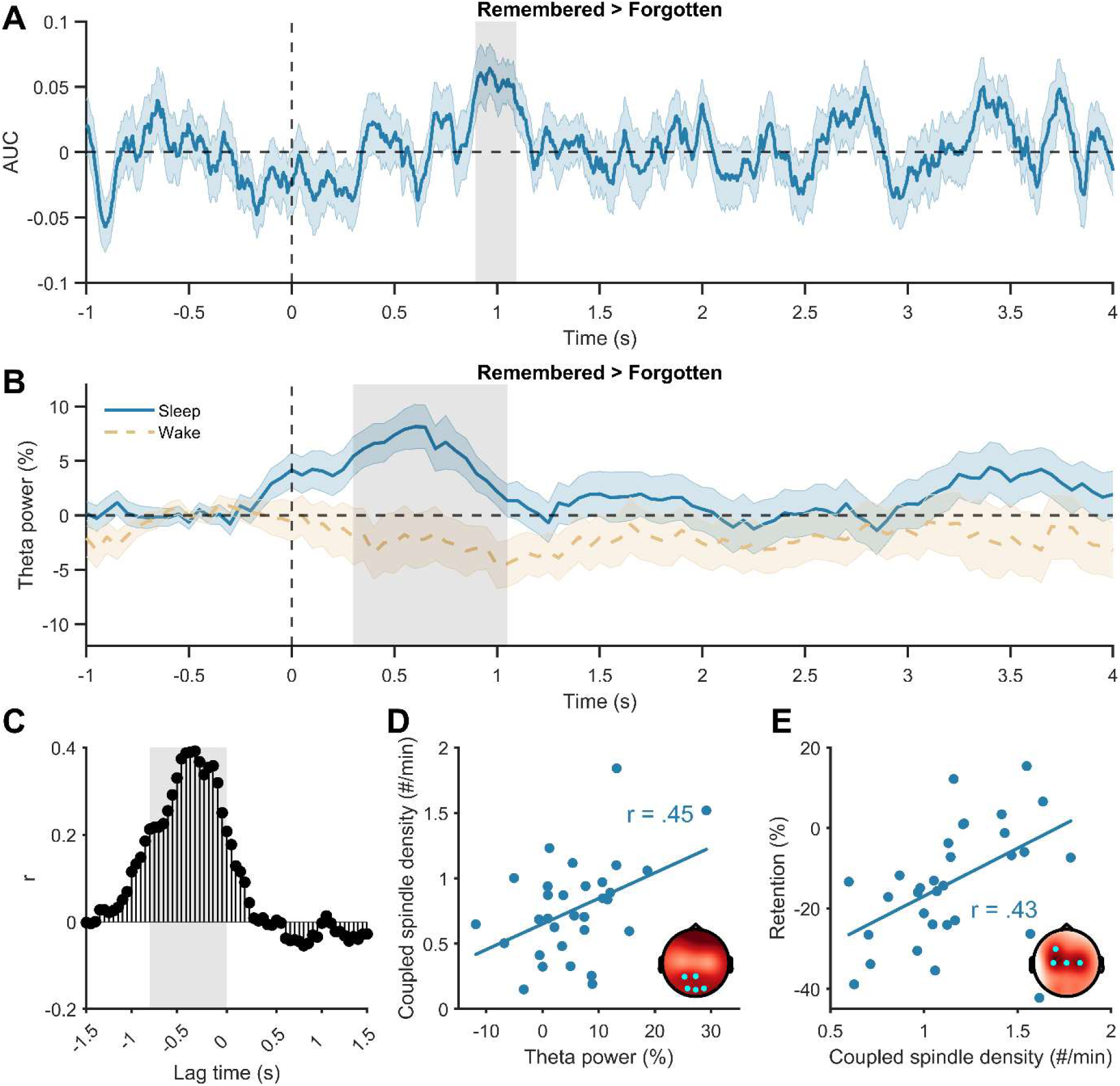
Encoding patterns differentially predict memory retention over a sleep or wake delay. **A** – Multivariate classification of EEG data at learning based on items that were subsequently remembered after sleep or wake. Significant classification above zero (extent of the cluster that is highlighted in grey, *p* < .05 corrected) indicates the time points of successful decoding. **B** – Theta power at encoding (normalised within-subject and session as a % change relative to the pre-stimulus baseline) was significantly higher for items successfully recalled after sleep compared to wake (significant time points in the cluster are highlighted in grey, *p* < .05 corrected). **C** – Cross-correlation between theta power at encoding (sleep > wake) and the multivariate classifier time series. Significant correlations (highlighted in grey, false discovery rate adjusted) at negative lags indicate that increased theta power at learning in the sleep condition predicted successful multivariate classification 0.45s later. **D** – Theta power at learning prior to sleep predicted a higher density of SO-coupled sleep spindles during subsequent sleep. The topographical inset highlights significant electrodes in the cluster (*p* < .05 corrected). **E –** Higher SO-coupled spindle density during sleep predicted better memory retention after sleep. The topography inset highlights significant electrodes in the cluster (*p* < .05 corrected).

### Theta oscillations uniquely predict memory retention over sleep

If theta oscillations underpin the tagging of memories at learning for consolidation during later sleep, then theta activity during learning should be predictive of memory performance after sleep, but not after wakefulness. To test this prediction, we decomposed the EEG data at learning into time-frequency representations (TFRs) and contrasted theta (3-8Hz) power for subsequently remembered > forgotten trials, separately for the sleep and wake conditions.

Theta power during successful learning was significantly higher for word-object pairs remembered after sleep than word-object pairs remembered after wake (0.3 – 1.05s, *t_sum_* = 42.53, *p_cluster_* = .032, *d_cluster_* = 0.54; **Figure 2B**). The increase in theta power was significantly greater than zero in the sleep condition (5.68% ± 8.09%; t (30) = 3.90, p < .001, d = 0.70), but not in the wake condition (-2.73% ± 12.18%; t (30) = -1.25, p = .22, d = -0.22). To confirm that our results were unique to the theta band, a broadband exploratory analysis was performed from 2-30Hz, with no additional significant clusters emerging (**Figure S3** for full TFRs and topographic effects, and **Figure S4** showing no difference in the event-related potentials). No difference in learning-related theta power was observed between the sleep and wake conditions for trials successfully recalled at the immediate test (no clusters formed; **Figure S2**). Taken together, these findings are consistent with the view that theta rhythms underpin the selective tagging of memories for consolidation during later sleep.

Interestingly, the peak in learning-related theta power for memories retained after sleep emerged ∼0.4s before the peak in multivariate classification accuracy for memories retained after sleep (theta power peak = 0.6s; **Figure 2B**, classifier fidelity peak = 0.98s; **Figure 2A**). This could imply that an early theta surge at learning triggers downstream processes that tag memories for consolidation during later sleep. To test for this, we adopted a cross-correlation approach which determined whether an early increase in theta power predicted later classification accuracy for memories retained after sleep. Consistent with this view, significant correlations were observed at lags ranging from -0.96 to -0.05s, peaking at -0.45s (*r*_max_ = .43; **Figure 2C**), aligning with the time difference in theta power and classifier peaks.

### Theta oscillations at learning predict slow oscillation-spindle coupling activity during sleep

If theta rhythms at learning represent a tagging mechanism for sleep-dependent consolidation, learning-related theta activity should be related to established oscillatory markers of sleep-dependent memory processing [2], namely the coupling between SOs and spindles (SO-spindle coupling [23]). We therefore predicted that theta power during successful learning would correlate with the density of SO-spindle coupling during later sleep, with the SO-spindle density in turn correlating with behavioural performance after sleep.

SO-spindle coupling events were detected using automated detectors [21]. Across all participants and electrodes, 16% ± 4% of spindles were coupled to an SO, far exceeding what would be expected by chance (**Supplementary Results**). We observed significant non-uniformity in the preferred phase of the SO to which spindles were coupled at 13/14 electrodes (*Z*s > 3.49, *p*s < .03). Sleep spindles preferentially coupled close to the positive peak of the SO (M = 20.96° ± 58.05°), in line with previous reports [21].

A robust linear regression model revealed a significant correlation between theta power during successful learning (i.e., for word-object pairs remembered after sleep) and SO-coupled spindle density (5 electrodes, *t_sum_* = 11.82, *p_cluster_* = .028, *r_cluster_* = .45; **Figure 2D**). In turn, memory retention across sleep was correlated with SO-coupled spindle density (4 electrodes, *t_sum_* = 10.62, *p_cluster_* = .042, *r_cluster_* = .43; **Figure 2E**).

No significant cluster emerged between theta power at learning and the density of spindles that were not coupled to SOs (i.e., uncoupled spindles) during sleep, nor between uncoupled spindle density and memory performance after sleep. Indeed, a multiple robust linear regression (Adjusted R^2^ = .21, *p* = .01) showed that coupled spindle density uniquely predicted memory retention over sleep (B [95%CI] = 25.6 [8.63, 42.57], *p* = .004), independent of uncoupled spindle density (B [95% CI] = -0.75 [-4.16, 2.66], *p* = .65). These results support the hypothesis that theta rhythms present at learning support the tagging of memories for sleep-dependent consolidation, which in turn drives the engagement of SO-coupled spindles during later sleep, resulting in selective memory strengthening and stabilisation.

Sleep spindles (and their coupling to SOs) are known to be trait-like [30] leaving open the possibility that the correlation between theta power, SO-coupled spindles and retention reflects more generalized individual differences rather than a specific mnemonic tagging process. However, theta power at learning for word-object pairs remembered after wakefulness was not significantly correlated with SO-spindle coupling density in the sleep condition. Moreover, no significant correlation was observed between SO-spindle coupling density in the sleep condition and retention in the wake condition. This demonstrates that the reported effects correspond to a unique, sleep-specific tagging mechanism and not trait-like oscillatory interactions.

## Discussion

Memory tags established at learning are thought to direct consolidation processes in later sleep [1–3], but neurobiological evidence for such a tagging mechanism in humans has yet to be established. Our results address this gap in three ways. First, we could reliably differentiate brain activity patterns at learning for memories that were subsequently retained over sleep or wakefulness, consistent with a dynamic tagging of memory for sleep-dependent memory processing. Second, theta activity at learning was uniquely predictive of memory retention after sleep and not wake, highlighting theta oscillations as a candidate mechanism through which sleep-specific memory tags are generated at encoding. Third, the magnitude of theta activity during pre-sleep learning was correlated with the density of SO-spindle coupling during subsequent sleep, which in turn was correlated with memory performance in the post-sleep retrieval test. Taken together, these findings demonstrate that theta-modulated tags acquired at learning act as an instructional cue for selective, overnight memory processing.

Our findings extend recent findings in the animal literature to humans and outline a potential mechanism that triages experiences for sleep-dependent memory processing. Interestingly, whereas studies in rodents have suggested reward or novelty cues generate tags for selective memory processing in later sleep [15,16,31], the word-object memories encoded by participants in the present study were matched for novelty and affective salience (all stimuli were familiar to participants and emotionally neutral). This approach was intentional as it allowed us to isolate generalised neurocognitive markers of mnemonic tagging. Yet, it prevented us from determining which stimulus properties increase the accessibility of memories to overnight memory processing. Given that the benefits of sleep for memory are often amplified for stimuli that are emotionally aversive [4–6], linked to reward [7,8] or deemed relevant for future engagements [9–11], an obvious next step will be to determine the impacts of these variables on our ability to decode tagging operations at learning in humans.

Theta oscillations have long been implicated in episodic learning and were uniquely associated with the tagging of memories for sleep-dependent consolidation in our current data. This builds on work in rodents showing that theta oscillations facilitate the late long-term potentiation necessary for selective memory strengthening [32,33], and research in humans demonstrating that theta activity at encoding maps onto the affective salience or distinctiveness of newly formed memories [19,20]. Mechanistically, theta oscillations are thought to support successful episodic learning by binding disparate aspects of an experience into a single, coherent representation [34,35]. Behavioural evidence has shown that the formation of integrated associations between previously unrelated items at encoding is key to observing a memory benefit of sleep [36–38]. Alongside this existing work, our findings might thus imply that theta-modulated, episodic binding is necessary for mnemonic tagging at learning and targeted processing during later sleep, with the propensity for such episodic binding influenced by the salience of the to-be-learned information.

In further support of the idea that theta-generated tags act as an instructional cue for selective overnight memory processing, theta activity during successful learning was correlated with SO-spindle coupling density during subsequent sleep, which in turn predicted the behavioural benefits of sleep for memory. Indeed, the interplay between global slow oscillations, thalamocortical spindles and hippocampal ripples is considered central to the reactivation of waking experience and integration of memories into long-term storage [39,40]. Our findings build on this prior work by demonstrating that these coordinated brain rhythms of deep, non-rapid eye movement sleep contribute to the selective strengthening of memories that have obtained theta-modulated tags during the initial learning phase. More broadly, this highlights how adaptive memory systems rely on interactions between the waking and sleeping brain.

While our data suggest that memory tags established at learning steer consolidation in the first period of sleep after learning, it remains unclear how they affect mnemonic operations across subsequent periods (i.e., nights) of sleep. Previous studies using targeted memory reactivation (TMR) protocols to bias sleep-dependent consolidation with learning-related sounds or odours, have shown that the effects of such interventions can take days or even weeks to emerge [41,42], meaning that tags formed at learning may likewise require multiple bouts of sleep to shape memory effectively. Furthermore, given recent views on the role of sleep in active forgetting [43], it is likewise possible that mnemonic tags established at learning may precipitate a targeted strengthening and *weakening* of prior experience to ensure that our memories remain aligned with our cognitive and emotional goals.

In sum, our findings provide evidence for a neurocognitive tagging signal in humans that predisposes memories for selective strengthening in later sleep, analogous to existing neurobiological data in the animal literature. This tagging process is driven by theta rhythms at learning, which act as an instructional cue to memory processes in the sleeping brain mediated by the interplay between SOs and sleep spindles. This serves as a mechanism through which the brain can retain salient experiences, helping us to quickly recall goal-relevant memories in an adaptive fashion to help guide our interactions with the world.

## Materials and Methods

### Participants

A total of 44 human participants enrolled in the study, of which 38 met inclusion criteria. Participants had no self-reported history of neurological, psychiatric, or sleep-related disorders, had normal or corrected-to-normal vision, were fluent in English, and performed adequately during a preliminary screening session (see **Procedure: Preliminary session**). Seven participants were excluded from the final analysis due to either not sleeping during their nap (n = 3) or being at floor or ceiling on at least one memory test (n = 4). Therefore, 31 participants were entered into the final analysis. Participants took part in exchange for an £80 e-voucher or course credit. The study was approved by the University of York Department Ethics Committee and written informed consent was obtained from all participants.

### Stimuli

A set of 480 words and 320 images served as experimental stimuli. Words were all concrete nouns describing natural objects (e.g., plants, animals, foodstuffs). Images were all of manmade objects presented on a plain white background, and were taken from the Bank of Standardized Stimuli (BOSS) database [44].

### Procedure

#### Preliminary session

Participants completed a preliminary visit to assess their eligibility for the study. Following informed consent, participants completed questionnaires assessing their subjective sleep quality (Pittsburgh Sleep Quality Index [45]), morning-evening preference (Morningness-eveningness questionnaire [46]) and daytime sleepiness (Epworth Sleepiness Scale [47]). They then performed an initial memory assessment, where they learned associations between 84 semantically unrelated word pairs [36,37]. Each word pair was presented for 5 s, and participants were instructed to form a vivid mental image combining the two word-pair referents together in a scene. Immediately after learning, memory for all word pairs was tested via a cued recall procedure. The first word was presented on the screen, and participants had 10 s to respond with the second word in the pair. Participants scoring between 30% and 80% were invited back for the main experiment. This was done to minimise the risk of participants performing at either floor or ceiling in the main study [48].

### Experimental sessions

#### Overview

The study consisted of two experimental sessions (sleep and wake condition) spaced 5 to 14 days parts. The order of the two conditions was counterbalanced across participants. Participants arrived at the sleep laboratory between 11am and 1pm and completed a retrospective sleep diary for the previous three nights to assess sleep quality in the nights leading up to the experimental session. There were no differences in self-reported sleep quality in the nights prior to the sleep or wake visit (**Table S1**). Next, participants were wired-up for electroencephalography (EEG) recordings. Following EEG setup, subjective alertness levels were assessed using the Stanford Sleepiness Scale (SSS) [49]. Next, participants were familiarized with the images before learning the word-object pairs (see task details below) and performing an immediate recall test. Next participants either took a nap (2 h sleep opportunity; sleep condition), or remained awake in the lab for 2 h, watching nature documentaries (wake condition). Afterwards, subjective alertness was assessed again using the SSS, before memory performance was tested for a second time (delayed recall test). There were no differences in self-reported alertness levels at either assessment (**Table S1**). All experimental tasks were presented using Psychtoolbox 3 [50] in MATLAB (MathWorks, Natick, MA, USA). Across all experimental tasks, stimulus presentation order was randomized across participants.

#### Familiarization

Participants were shown all of the to-be-encoded objects and their names/descriptors. This was done to facilitate subsequent word-object learning and to provide the proper object names for later cued recall. Each trial started with a fixation cross for 1 s (± 0.2 s jitter), followed by an object with its name presented above for 2 s. For each trial, participants were asked to think how often they would use or interact with the object in everyday life.

#### Learning

Participants learned 160 word-object pairs. On each trial, a fixation cross was presented for 1 s (± 0.2 s jitter), followed by one word-object pair for 4 s. Participants were told to visualize a scene that combined the referent of the word and the object into a single coherent scene (e.g., if the word COFFEE was paired with an image of a football, participants might visualize an image of a cup of coffee balanced on a football). This type of strategy has been shown to enhance sleep-dependent memory consolidation in previous work [36,37]. After 4 s, the word-object pair faded away from the screen and was replaced by an instruction to ‘Think’ for a further 4 s. During this period, participants were instructed to hold the mental image they had formed in their mind to facilitate successful learning [36]. At the end of the trial, participants were instructed to indicate the vividness of the mental image on a scale from 1 (no image formed) to 4 (very vivid visualization). Average visualisation vividness was 2.78 ± 0.38 and did not differ between sleep and wake conditions (*p* = .58). Each trial was shown once during the learning phase, and participants were told beforehand that their memory for the associations would be tested.

#### Recall

One half of the word-object pairs were tested at the immediate recall test and the other half at the delayed recall test. This permits an assessment of sleep vs wake on memory retention whist preventing retrieval practice effects that might emerge when testing the same items at the immediate and delayed assessments [28]. Other than that, the immediate and delayed recall tests were identical. Each test comprised 80 randomly chosen “old” words from the learning phase intermixed with 40 “new” words not seen by the participant during learning. Each trial began with a fixation cross (1 s ± 0.2 s jitter), after which a word was displayed on the screen for 4 s. During this time, participants were instructed to bring to mind the object originally paired with the word (if they recognized the word). After 4 s, participants indicated whether the word was “old” (i.e., seen during learning) or “new” (i.e., not seen during learning). A “don’t know” option was also presented. To minimize guessing, participants were instructed to only select “old” or “new” if they were confident in their judgement and to press the “don’t know” option when they were not confident. For each identified “old” trial, participants were then given 10 s to type a description of the object that was originally paired with the word.

#### Sleep delay

Participants were given a 2 h opportunity to nap in a laboratory bedroom whilst EEG was monitored. After waking up, participants were given at least 10 min to mitigate the effects of sleep inertia before continuing with the post-sleep memory test.

#### Wake delay

Participants remained in the sleep lab for 2 h watching nature documentaries. They were prohibited from other tasks such as reading or using their mobile phones.

#### Electroencephalography

EEG was acquired with either an Embla NDx or Embla N7000 system using RemLogic 3.4 software. For all participants, the same EEG device was used for both experimental sessions. Gold-plated electrodes were attached to the scalp for EEG (14 electrodes, positioned according to the 10-20 system), above the right eye and below the left eye for EOG, and on the left and right side of the chin for EMG. Ground and reference electrodes were placed above the left and right eyebrow (corresponding to Fp1 and Fp2 of the 10-20 system). Finally, two electrodes were positioned on the left and right mastoid for offline re-referencing (see EEG analysis below). Data were recorded at 256Hz (Embla NDx, later downsampled to 200 Hz) or 200Hz (Embla N7000), and impedances were kept below 5kΩ.

### Data analysis

#### Behaviour

To assess memory performance, we calculated the proportion of correctly recalled objects relative to the number of correctly recognized words (i.e., the number of hits). In other words, object memory was only considered when participants correctly recognized the word. An object was considered correctly recalled if either 1) the participant typed the same descriptor that was shown during the familiarization phase or 2) the description provided unambiguously matched the object. Correctly recalling the object when cued with the word indicates successful associative memory retrieval, which is the focus here. For completeness, item memory scores (hit and false alarm rates) are provided in **Table 1**. To quantify retention over the delay period, we calculated a relative change in object recall as [(delayed recall – immediate recall) / immediate recall]. This measure quantifies memory changes from the immediate to delayed test while controlling for immediate memory performance [36].

### EEG

#### Preprocessing – wake

EEG data were preprocessed using functions from the FieldTrip toolbox for MATLAB [51] and custom MATLAB scripts. A notch filter was applied at 50Hz to remove electrical line noise and a high-pass filter was applied at .3Hz to remove low frequency artefacts. EEG channels were then re-referenced to the average of the two mastoid channels, before being epoched from -2 to +6 s around stimulus pair onset during learning. Noisy channels and epochs were identified and removed via visual inspection (*ft_visreject* function in FieldTrip; see **Table S3** for trial counts) and bad channels were interpolated using a weighted average of the neighbouring channels. To correct for ocular artefacts, the epoched data were subjected to an independent component analysis, and ICA components reflecting eye blinks and movements were rejected.

#### Preprocessing – sleep

Nap data were first manually scored in 30-s epochs in accordance with AASM criteria [52]. Artefactual epochs were detected using an automated algorithm [53]. For each EEG channel, we calculated per-epoch summary metrics of three Hjorth parameters (signal activity, mobility, and complexity). Any epochs in which any one of these parameters was > 3 standard deviations from the mean, for one or more channels, were marked as artefact. Artefact detection was performed twice (in case of extreme outliers), and separately for each sleep stage (given inherent differences in the EEG signal between different sleep stages) [53]. Subsequently, all artefact free NREM (stage N2 + N3) epochs were retained for further analysis.

#### Multivariate analysis

We performed a linear classification of single-trial EEG data at learning using the MVPA-Light toolbox for MATLAB and an LDA classifier [54]. The classifier was trained to discriminate between learning trials for which the object memories were remembered in the sleep and wake conditions. Learning data were baseline corrected (-1 to 0 s relative to stimulus onset), linear trends were removed, and data were z-scored across all trials within each condition (sleep or wake) and for each timepoint separately. The two datasets were then smoothed using running average windows of 150 ms. The 14 EEG channels served as features and a different classifier was trained and tested on every sample from -1 to 4 s around stimulus onset. Area under the ROC curve was our classification metric, which indexes the mean accuracy with which a randomly chosen pair of learning trials could be assigned to their correct class (learning before sleep or wake; 0.5 = chance level performance). To avoid overfitting, data were split into training and test sets using a fivefold cross-validation. Because cross-validation results are stochastic due to the random assignment of trials to folds, the analysis was repeated ten times and the results averaged. To account for differences in the number of trials remembered after sleep compared to after wake, the number of trials per condition were equated by subsampling a random selection of trials from the higher trial count condition to match the condition with the fewer trials. This resulted in M = 32 (SD = 15) trials per condition. To assess whether classifier accuracy was significantly above chance, surrogate decoding performance was calculated by shuffling the training labels 250 times, and taking the average of these permutations [25]. This provides, for each participant, a baseline decoding performance that can be compared to the classifier run on the real class labels. Multivariate classification was performed on the preprocessed EEG time series, rather than the theta time-frequency power time series (see next section), so as not to filter out potentially informative information at other frequencies present in the multivariate signal.

An important caveat of our approach is that, by necessity, the sleep and wake conditions were performed on separate experimental days. Non-specific between-session differences, such as differences in impedance levels and the exact positioning of electrodes could thus be picked up by the classifier. It was therefore crucial to ensure that any signal discriminating between the sleep and wake conditions did not more parsimoniously reflect significant classification of the experimental sessions. To account for this, a set of control analyses were carried out. First, we ran the classifier again, but this time distinguishing trials that were *forgotten* after sleep compared to *forgotten* after wake (i.e., the Forgotten classifier). By comparing this classifier to the first Remember classifier, we could look for periods where classification accuracy was significantly higher for later remembered vs not remembered trials. If classification accuracy was higher for one vs the other, this would speak against more generic between visit differences, because we would not expect broad between-sessions differences to impact the memory conditions differently. As a second control analysis, we ran a third classifier to discriminate between Visit 1 and Visit 2 (the Visit classifier). This is dissociable from the sleep and wake condition because the assignment of visit order (sleep first or wake first) was counterbalanced across participants.

For our primary analysis, we performed a double subtraction of the classifier time series. First, we subtracted the Visit classifier time course from the Remember classifier and from the Forgotten classifier. Thus, any remaining values above zero would reflect genuine sleep vs wake condition differences for subsequently remembered and forgotten word-image pairings, having removed more generic between-visit factors. Second, we subtracted the visit-corrected Forgotten classifier time series from the visit-corrected Remember classifier time series. This resulted in a single time series where values above zero would reflect a remember-specific signal that differed depending on whether learning occurred prior to sleep or wake.

As a third control, we wanted to rule out any possibility that significant classification could be explained by participants simply engaging in a different cognitive strategy or being in a broadly different mental state (e.g., by knowing that they would be taking a nap or remaining awake across the delay). If this were the case, then significant classification between the sleep and wake conditions would also be possible on learning trials that were tested at the immediate, pre-delay test. Therefore, we repeated the above procedure but focused on learning trials that were remembered/forgotten at the immediate, pre-delay test. This control analysis was performed to confirm that effects were specific to testing after sleep.

#### Time-frequency analysis

Preprocessed data were convolved with a five-cycle Hanning taper (*ft_freqanalysis* function in FieldTrip, using the *mtmconvol* method) from -2 to 6 s relative to stimulus onset in steps of 50ms, and from 3 -30Hz in steps of 1Hz. To avoid edge artefacts, subsequent analyses focused on the -1 to 4 s time window (consistent with the classification analysis). Artefact rejection was performed on single trial time-frequency representations (TFRs) using a data-driven approach [48]: power values that exceeded the 85th percentile across all time/frequency points and trials were rejected and removed from subsequent analyses. TFRs were converted into a percent power change relative to a baseline interval of -0.4 s to -0.2 s before stimulus onset. This window was chosen to mitigate baseline contamination by post stimulus activity while preserving proximity to stimulus onset [48]. Theta power estimates were extracted by averaging power values between 3-8Hz. Then, the trial-averaged signal for subsequently forgotten word-object pairs was subtracted from the trial-averaged signal for subsequently remembered word-object pairs, separately for the sleep and wake delay conditions. Thus, the final TFR values entered into statistical analysis reflect theta power signatures of successful learning prior to a delay containing sleep or wake. To again confirm specificity to post-sleep testing, we repeated the above procedure but focusing on trials remembered/forgotten at the immediate, pre-delay test.

#### Slow oscillation-spindle event detection

First, each individual’s peak spindle frequency was identified through visual inspection of the NREM power spectrum. The largest, most prominent peak in the 12-16 Hz range was considered that individual’s spindle peak frequency. Spindles were then automatically detected using a wavelet based detector [21,55,56]. The raw EEG signal was convolved with a complex Morlet wavelet, with the wavelet peak frequency set at that individual’s spindle peak frequency, and the bandwidth of the wavelet (FWHM) set as a 1.3Hz range centered on the peak frequency [21,57]. A spindle was detected whenever the wavelet-filtered signal exceeded a threshold of 6 times the median signal amplitude for a minimum of 0.4s. The threshold of 6 times the median has been empirically determined to maximize between-class (spindle, non-spindle) variance in the wavelet-filtered signal in previous work using a nap paradigm and healthy young adults [21].

Slow oscillations were then detected using a second automated algorithm [58]. Data were initially bandpass filtered between 0.5-4Hz and all positive-to-negative zero crossings were identified. Candidate slow oscillations were marked if two such consecutive crossings fell 0.8-2 s apart (i.e., 0.5-1.25Hz, consistent with the SO frequency). Peak-to-peak amplitudes of all candidate oscillations were determined, and those in the top quartile (i.e., with the highest amplitudes) were retained as SOs. For SO-spindle coupling detection, the Hilbert transform was applied to extract the instantaneous phase of the SO signal and the instantaneous amplitude of the spindle signal. For each detected spindle, its peak amplitude was determined. If the spindle peak was found to occur during the time course of any detected 0.5 – 1.25Hz slow oscillations (i.e., between the two positive-to-negative crossings), the event was marked as a SO-spindle coupling event. Our primary measure of interest was the SO-spindle coupling density (i.e., the number of SO-coupled spindles per min of NREM sleep) [21,59,60].

#### Statistical analysis

Differences between the sleep and wake conditions for a) retention over the delay and b) immediate recall performance were assessed with paired-samples t-tests. For multivariate classification analyses, comparison of classifier accuracy against the surrogate baseline distribution was performed using a paired-samples t-test at each time point. Analysis of the classifier time series following the double subtraction was achieved via a 1-sample t-test against zero at every time point. To examine sleep-wake differences in TFRs, between-condition differences were compared by way of a paired-samples t-test conducted at every time point. For all analyses, multiple comparisons across timepoints/electrodes, were controlled for using a cluster-based permutation method [61]. A cluster-corrected p < .05 was deemed statistically significant. Effect sizes were derived by averaging across all significant time/electrode points that contributed to the cluster, expressed as Cohen’s d.

To test for time-lagged associations between TFR power and multivariate classification, the cross-correlation between the TFR time series and lagged copies of the classifier time series (from -1.5 to 1.5s lags, in 50ms intervals) were computed. Here, a negative lag would indicate that theta activity predicts later multivariate classification, whereas a positive lag would suggest the opposite. Multiple comparisons across multiple lags in the cross-correlation analysis were controlled using the False Discovery Rate [62]. Correlation analyses between theta power at learning, coupled spindle density (number of coupled spindles per min), and memory retention were performed as robust linear regressions to minimise the influence of outliers. Multiple comparisons across electrodes were controlled using cluster-based permutation testing. Pearson’s *r* values are reported to quantify effect size.

## Acknowledgments

We are grateful to members of the Sleep, Language, and Memory (SLAM) lab at the University of York and the Staresina Lab at the University of Oxford for fruitful discussions of the data.

## Supplementary Materials

### Supplementary Results

To determine whether the co-occurrence of SOs and spindles reflects a “true” coupling of the two oscillations, we needed to ensure that the number of coupling events exceeded what would be expected by chance, given the amount of time spent in non-rapid eye movement sleep, and the number of detected slow oscillations and spindles. To this end, the observed SO and spindle signal was compared to a randomized signal where the spindle signal was circularly shifted, and coupling to SOs was re-calculated [1, 2]. The start of the shifted signal is set to a random position, and early entries are sequentially moved to after the end of the original signal. By shifting the spindle signal in this fashion, the temporal relationship between SOs and spindles is disrupted, whilst retaining the signal properties of the original signal (i.e., the distribution of SOs and spindles). This was performed over 1000 iterations to generate a null distribution of SO-spindle coupling. The null distribution was then compared with the observed number of coupling events. Across each participant/electrode, we calculated the percentage of participants and electrodes where the degree of observed coupling was significantly higher (p < .05) than what would be expected by chance. Across all electrodes, 29/31 (94%) of participants exhibited a rate of coupling that significantly exceeded what would be expected by chance (binomial test, p < .001).

---

**Table S1.** Participant demographics and questionnaire measures.

|  | M | SD |  |  |  |  |
| --- | --- | --- | --- | --- | --- | --- |
| Age (years) | 20 | 1 |  |  |  |  |
| Sex | N | % |  |  |  |  |
| Female | 21 | 68 |  |  |  |  |
| Male | 10 | 32 |  |  |  |  |
|  | M | SD |  |  |  |  |
| PSQI | 4.45 | 1.95 |  |  |  |  |
| MEQ | 48.7 | 7.38 |  |  |  |  |
| ESS | 5.23 | 2.88 |  |  |  |  |
|  | Sleep visit |  | Wake visit |  | *t* | *p* |
|  | M | SD | M | SD |  |  |
| <u>Sleep diary</u> |  |  |  |  |  |  |
| TST (h) | 8 | 2 | 8 | 2 | 0.33 | .75 |
| SOL (min) | 20 | 22 | 21 | 29 | 1.28 | .21 |
| WASO (min) | 7 | 8 | 8 | 10 | 0.90 | .38 |
| SE (%) | 95 | 4 | 94 | 5 | 0.82 | .42 |
| <u>Stanford sleepiness scale</u> |  |  |  |  |  |  |
| Learning | 2.19 | 0.83 | 2.03 | 0.84 | 0.41 | .69 |
| Recall 2 | 2.81 | 1.01 | 2.71 | 1.19 | 0.33 | .75 |

*Note.* M = Mean, SD = standard deviation. PSQI = Pittsburgh sleep quality index global score. A higher score indicates worse subjective sleep quality (theoretical range: 0-21) MEQ = Morningness-eveningness questionnaire. A higher score indicates greater preference for mornings (theoretical range: 16-86) ESS = Epworth sleepiness scale. A higher score indicates higher subjective daytime sleepiness (theoretical range: 0-24). TST = total sleep time, SOL = sleep onset latency, WASO = Wake after sleep onset, SE = Sleep efficiency (% of time in bed that was spent asleep). Sleep diary measures calculated as the average of the three nights prior to the experimental day.

**Table S2.** Sleep architecture.

|  | M | SD |
| --- | --- | --- |
| Total sleep time (min) | 84 | 31 |
| N1 (%) | 13.52 | 8.25 |
| N2 (%) | 54.78 | 12.23 |
| N3 (%) | 19.98 | 14.59 |
| REM (%) | 11.71 | 12.07 |
| Sleep spindles (N) | 367 | 178 |
| Slow oscillations (N) | 303 | 134 |
| SO-spindle coupling events (N) | 57 | 28 |

*Note.* M = mean, SD = standard deviation. SO = slow oscillation. Spindle, SO, and SO-spindle coupling counts derived from electrode Cz.

**Table S3.** Trial counts for EEG analyses.

|  | Immediate recall |  | Delayed recall |  |
| --- | --- | --- | --- | --- |
|  | Remembered | Forgotten | Remembered | Forgotten |
|  | M (SD) | M (SD) | M (SD) | M (SD) |
| Sleep | 47 (16) | 38 (20) | 38 (18) | 36 (18) |
| Wake | 48 (15) | 38 (22) | 32 (15) | 41 (15) |

*Note.* M = mean, SD = standard deviation. Average trial counts presented for information only. Note that for all analyses, trial-counts were matched per-participant by randomly selecting a subset of trials from the higher count condition to match the lowest trial condition.

**Figure S1.**
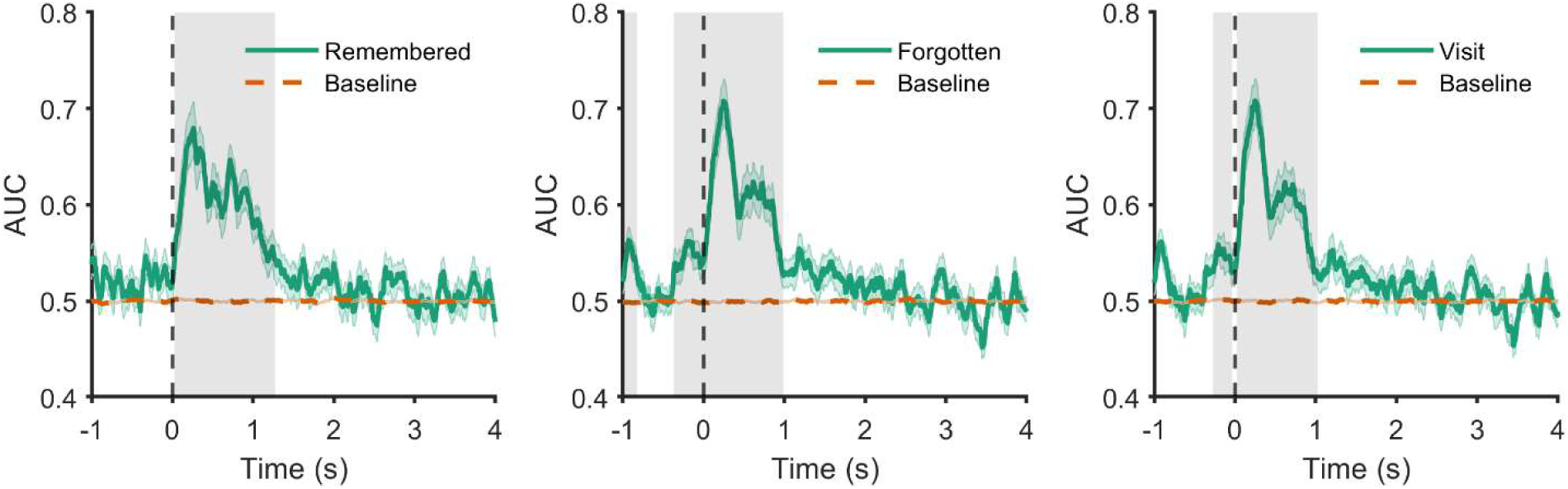
Classifier performance vs baseline. Ability of classifier to decode learning trials into subsequently remembered across the sleep or wake delay (left); subsequently forgotten across the sleep or wake delay (middle); or decoding of experimental visit independent of delay condition or memory status (right). Classifier accuracy was compared to a surrogate decoding baseline, which was estimated by shuffling the training labels 250 times. Significant clusters (i.e. where classification exceeded chance levels) are highlighted in grey. Shaded areas around the lines indicates the standard error of the mean.

**Figure S2.**
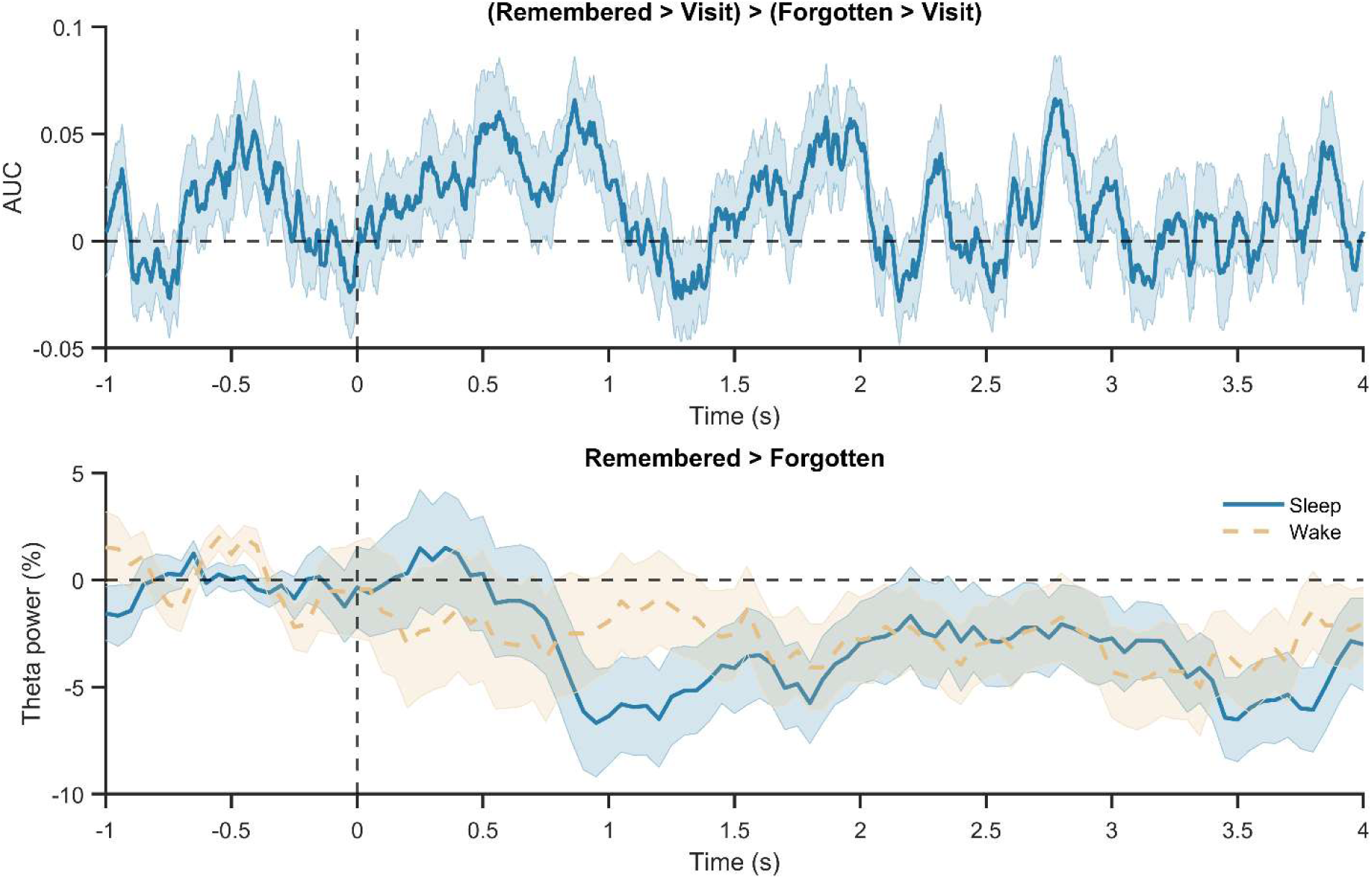
Top. Multivariate classification could not classify delay type (sleep or wake) based on learning trials that were remembered at the immediate test. Bottom – No theta power differences at learning for trials recalled at the immediate test prior to the sleep or wake delay.

**Figure S3.**
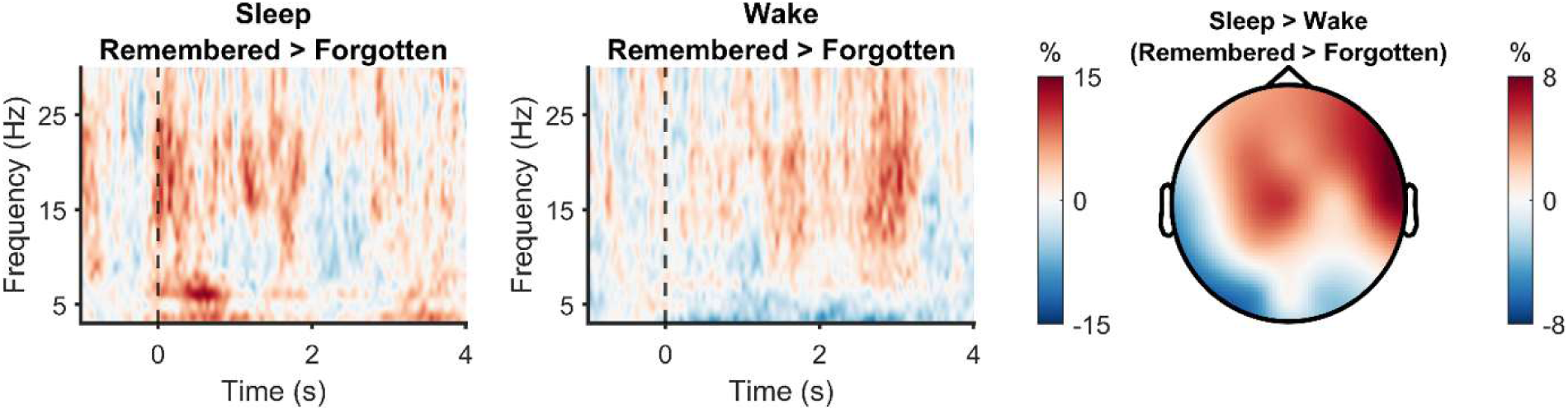
Time-frequency representations at learning prior to sleep (left) and wake (middle). Warmer colours indicate a greater % increase (relative to pre-stimulus baseline) in spectral power for remembered compared to forgotten trials. The right-hand plot depicts the spatial extent of theta-related (3-8Hz) learning activity for word-object pairs that were remembered (> forgotten) after sleep (> wake).

**Figure S4.**
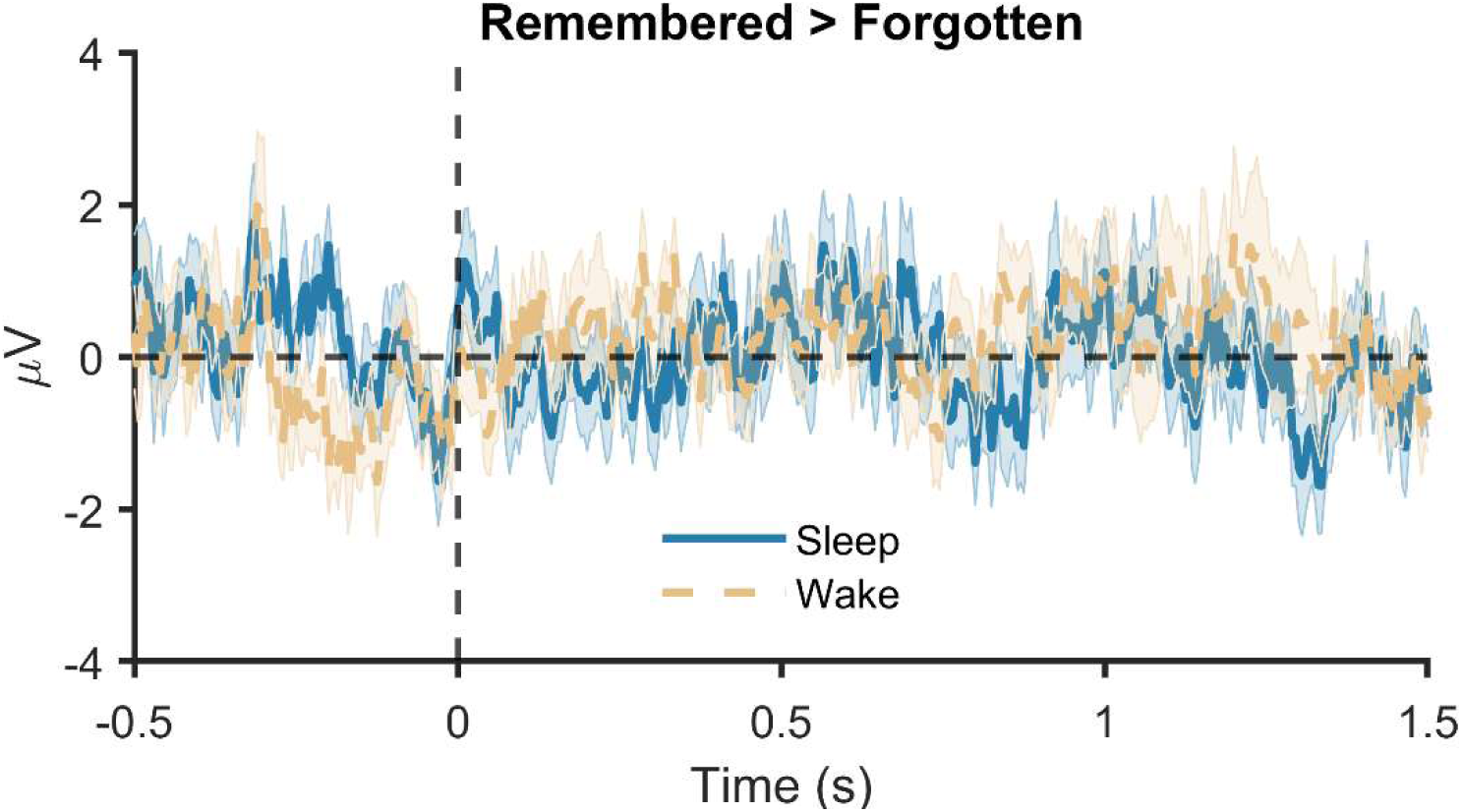
Event-related potentials during learning. No difference in ERP activity between the sleep and wake conditions were observed. Shaded area around the line indicates the standard error.

## References

1. Stickgold R, Walker MP. Sleep-dependent memory triage: evolving generalization through selective processing. Nat Neurosci. 2013;16: 139–145. doi:10.1038/nn.3303

2. Cowan ET, Schapiro AC, Dunsmoor JE, Murty VP. Memory consolidation as an adaptive process. Psychon Bull Rev. 2021;28: 1796–1810. doi:10.3758/s13423-021-01978-x

3. Born J, Wilhelm I. System consolidation of memory during sleep. Psychol Res. 2012;76: 192–203. doi:10.1007/s00426-011-0335-6

4. Payne JD, Stickgold R, Swanberg K, Kensinger EA. Sleep preferentially enhances memory for emotional components of scenes. Psychol Sci. 2008;19: 781–788. doi:10.1111/j.1467-9280.2008.02157.x

5. Cox R, van Bronkhorst MLV, Bayda M, Gomillion H, Cho E, Parr ME, et al. Sleep selectively stabilizes contextual aspects of negative memories. Sci Rep. 2018;8: 17861. doi:10.1038/s41598-018-35999-9

6. Denis D, Sanders KEG, Kensinger EA, Payne JD. Sleep preferentially consolidates negative aspects of human memory: Well-powered evidence from two large online experiments. Proceedings of the National Academy of Sciences. 2022;119: e2202657119. doi:10.1073/pnas.2202657119

7. Fischer S, Born J. Anticipated reward enhances offline learning during sleep. J Exp Psychol Learn Mem Cogn. 2009;35: 1586–1593. doi:10.1037/a0017256

8. Igloi K, Gaggioni G, Sterpenich V, Schwartz S. A nap to recap or how reward regulates hippocampal-prefrontal memory networks during daytime sleep in humans. eLife. 2015;4: e07903. doi:10.7554/eLife.07903

9. Wilhelm I, Diekelmann S, Molzow I, Ayoub A, Mölle M, Born J. Sleep selectively enhances memory expected to be of future relevance. J Neurosci. 2011;31: 1563–1569. doi:10.1523/JNEUROSCI.3575-10.2011

10. Cunningham TJ, Bottary R, Denis D, Payne JD. Sleep spectral power correlates of prospective memory maintenance. Learn Mem. 2021;28: 291–299. doi:10.1101/lm.053412.121

11. Scullin MK, McDaniel MA. Remembering to Execute a Goal: Sleep on It! Psychological Science. 2010;21: 1028–1035. doi:10.1177/0956797610373373

12. Davidson P, Jönsson P, Carlsson I, Pace-Schott E. Does Sleep Selectively Strengthen Certain Memories Over Others Based on Emotion and Perceived Future Relevance? NSS. 2021;13: 1257–1306. doi:10.2147/NSS.S286701

13. Saletin JM, Walker MP. Nocturnal mnemonics: sleep and hippocampal memory processing. Front Neurol. 2012;3: 59. doi:10.3389/fneur.2012.00059

14. Kim SY, Payne JD. Neural correlates of sleep, stress, and selective memory consolidation. Current Opinion in Behavioral Sciences. 2020;33: 57–64. doi:10.1016/j.cobeha.2019.12.009

15. Huelin Gorriz M, Takigawa M, Bendor D. The role of experience in prioritizing hippocampal replay. Nat Commun. 2023;14: 8157. doi:10.1038/s41467-023-43939-z

16. Yang W, Sun C, Huszár R, Hainmueller T, Kiselev K, Buzsáki G. Selection of experience for memory by hippocampal sharp wave ripples. Science. 2024;383: 1478–1483. doi:10.1126/science.adk8261

17. King J-R, Dehaene S. Characterizing the dynamics of mental representations: the temporal generalization method. Trends Cogn Sci (Regul Ed). 2014;18: 203–210. doi:10.1016/j.tics.2014.01.002

18. Heib DPJ, Hoedlmoser K, Anderer P, Gruber G, Zeitlhofer J, Schabus M. Oscillatory theta activity during memory formation and its impact on overnight consolidation: a missing link? J Cogn Neurosci. 2015;27: 1648–1658. doi:10.1162/jocn_a_00804

19. Gruber MJ, Watrous AJ, Ekstrom AD, Ranganath C, Otten LJ. Expected reward modulates encoding-related theta activity before an event. Neuroimage. 2013;64: 68–74. doi:10.1016/j.neuroimage.2012.07.064

20. Heinbockel H, Quaedflieg CWEM, Schneider TR, Engel AK, Schwabe L. Stress enhances emotional memory-related theta oscillations in the medial temporal lobe. Neurobiology of Stress. 2021;15: 100383. doi:10.1016/j.ynstr.2021.100383

21. Denis D, Mylonas D, Poskanzer C, Bursal V, Payne JD, Stickgold R. Sleep Spindles Preferentially Consolidate Weakly Encoded Memories. J Neurosci. 2021;41: 4088–4099. doi:10.1523/JNEUROSCI.0818-20.2021

22. Staresina BP. Coupled sleep rhythms for memory consolidation. Trends in Cognitive Sciences. 2024;28: 339–351. doi:10.1016/j.tics.2024.02.002

23. Denis D, Cairney SA. Electrophysiological Mechanisms of Memory Consolidation in Human Non-rapid Eye Movement Sleep. Curr Sleep Medicine Rep. 2024;10: 181–190. doi:10.1007/s40675-024-00291-y

24. Ng T, Noh E, Spencer RM. Does slow oscillation-spindle coupling contribute to sleep-dependent memory consolidation? A Bayesian meta-analysis. eLife. 2024;13. doi:10.7554/eLife.101992.1

25. Schreiner T, Petzka M, Staudigl T, Staresina BP. Endogenous memory reactivation during sleep in humans is clocked by slow oscillation-spindle complexes. Nature Communications. 2021;12: 3112. doi:10.1038/s41467-021-23520-2

26. Studte S, Bridger E, Mecklinger A. Sleep spindles during a nap correlate with post sleep memory performance for highly rewarded word-pairs. Brain Lang. 2017;167: 28–35. doi:10.1016/j.bandl.2016.03.003

27. Denis D, DiPietro C, Spreng RN, Schacter DL, Stickgold R, Payne JD. Sleep and retrieval practice both strengthen and distort story recollection. SLEEP Advances. 2024;5: zpae083. doi:10.1093/sleepadvances/zpae083

28. Bäuml K-HT, Holterman C, Abel M. Sleep can reduce the testing effect: It enhances recall of restudied items but can leave recall of retrieved items unaffected. Journal of Experimental Psychology: Learning, Memory, and Cognition. 2014;40: 1568–1581. doi:10.1037/xlm0000025

29. Paller KA, Wagner AD. Observing the transformation of experience into memory. Trends in Cognitive Sciences. 2002;6: 93–102. doi:10.1016/S1364-6613(00)01845-3

30. Cox R, Mylonas DS, Manoach DS, Stickgold R. Large-scale structure and individual fingerprints of locally coupled sleep oscillations. Sleep. 2018;41: zsy175. doi:10.1093/sleep/zsy175

31. Lansink CS, Goltstein PM, Lankelma JV, McNaughton BL, Pennartz CMA. Hippocampus Leads Ventral Striatum in Replay of Place-Reward Information. PLOS Biology. 2009;7: e1000173. doi:10.1371/journal.pbio.1000173

32. Leung LS, Law CSH. Phasic modulation of hippocampal synaptic plasticity by theta rhythm. Behavioral Neuroscience. 2020;134: 595–612. doi:10.1037/bne0000354

33. Cao G, Harris KM. Augmenting saturated LTP by broadly spaced episodes of theta-burst stimulation in hippocampal area CA1 of adult rats and mice. Journal of Neurophysiology. 2014;112: 1916–1924. doi:10.1152/jn.00297.2014

34. Griffiths BJ, Martín-Buro MC, Staresina BP, Hanslmayr S. Disentangling neocortical alpha/beta and hippocampal theta/gamma oscillations in human episodic memory formation. NeuroImage. 2021;242: 118454. doi:10.1016/j.neuroimage.2021.118454

35. Hanslmayr S, Staresina BP, Bowman H. Oscillations and Episodic Memory: Addressing the Synchronization/Desynchronization Conundrum. Trends in Neurosciences. 2016;39: 16–25. doi:10.1016/j.tins.2015.11.004

36. Denis D, Schapiro AC, Poskanzer C, Bursal V, Charon L, Morgan A, et al. The roles of item exposure and visualization success in the consolidation of memories across wake and sleep. Learn Mem. 2020;27: 451–456. doi:10.1101/lm.051383.120

37. Denis D, Bottary R, Cunningham TJ, Tcheukado M-C, Payne JD. The influence of encoding strategy on associative memory consolidation across wake and sleep. Learn Mem. 2023;30: 185–191. doi:10.1101/lm.053765.123

38. Ashton JE, Staresina BP, Cairney SA. Sleep bolsters schematically incongruent memories. PLOS ONE. 2022;17: e0269439. doi:10.1371/journal.pone.0269439

39. Denis D, Cairney SA. Neural reactivation during human sleep. Emerging Topics in Life Sciences. 2023;7: 487–498. doi:10.1042/ETLS20230109

40. Klinzing JG, Niethard N, Born J. Mechanisms of systems memory consolidation during sleep. Nat Neurosci. 2019;22: 1598–1610. doi:10.1038/s41593-019-0467-3

41. Cairney SA, Guttesen AÁV, El Marj N, Staresina BP. Memory Consolidation Is Linked to Spindle-Mediated Information Processing during Sleep. Curr Biol. 2018;28: 948–954. doi:10.1016/j.cub.2018.01.087

42. Rakowska M, Abdellahi MEA, Bagrowska P, Navarrete M, Lewis PA. Long term effects of cueing procedural memory reactivation during NREM sleep. NeuroImage. 2021;244: 118573. doi:10.1016/j.neuroimage.2021.118573

43. Cairney SA, Horner AJ. Forgetting unwanted memories in sleep. Trends in Cognitive Sciences. 2024;28: 881–883. doi:10.1016/j.tics.2024.07.011

44. Brodeur MB, Dionne-Dostie E, Montreuil T, Lepage M. The Bank of Standardized Stimuli (BOSS), a New Set of 480 Normative Photos of Objects to Be Used as Visual Stimuli in Cognitive Research. PLOS ONE. 2010;5: e10773. doi:10.1371/journal.pone.0010773

45. Buysse DJ, Reynolds C, Monk T, Berman S, Kupfer D. The Pittsburgh Sleep Quality Index (PSQI): A new instrument for psychiatric practice and research. Psychiatry Research. 1989;28: 193–213. doi:10.1016/0165-1781(89)90047-4

46. Horne JA, Ostberg O. A self-assessment questionnaire to determine morningness-eveningness in human circadian rhythms. Int J Chronobiol. 1976;4: 97–110.

47. Johns MW. A new method for measuring daytime sleepiness: the Epworth sleepiness scale. Sleep. 1991;14: 540–545. doi:10.1093/sleep/14.6.540

48. Guttesen A á V, Gaskell MG, Madden EV, Appleby G, Cross ZR, Cairney SA. Sleep loss disrupts the neural signature of successful learning. Cerebral Cortex. 2022; bhac159. doi:10.1093/cercor/bhac159

49. Hoddes E, Dement W, Zarcone V. The development and use of the Stanford sleepiness scale. Psychophysiology. 1972;9: 150.

50. Kleiner M, Brainard D, Pelli D, Ingling A, Murray R, Broussard C. What’s new in psychtoolbox-3. Perception. 2007;36: 1–16.

51. Oostenveld R, Fries P, Maris E, Schoffelen J-M. FieldTrip: Open source software for advanced analysis of MEG, EEG, and invasive electrophysiological data. Comput Intell Neurosci. 2011;2011: 156869. doi:10.1155/2011/156869

52. Iber C, Ancoli-Israel S, Chesson A, Quan SF. The AASM Manual for the Scoring of Sleep and Associated Events: Rules, Terminology and Technical Specification. Westchester, IL: Americian Academy of Sleep Medicine; 2007.

53. Purcell SM, Manoach DS, Demanuele C, Cade BE, Mariani S, Cox R, et al. Characterizing sleep spindles in 11,630 individuals from the National Sleep Research Resource. Nat Commun. 2017;8: 15930. doi:10.1038/ncomms15930

54. Treder MS. MVPA-Light: A Classification and Regression Toolbox for Multi-Dimensional Data. Frontiers in Neuroscience. 2020;14: 289. doi:10.3389/fnins.2020.00289

55. Mylonas D, Tocci C, Coon WG, Baran B, Kohnke EJ, Zhu L, et al. Naps reliably estimate nocturnal sleep spindle density in health and schizophrenia. Journal of Sleep Research. 2019;29: e12968. doi:10.1111/jsr.12968

56. Wamsley EJ, Tucker MA, Shinn AK, Ono KE, McKinley SK, Ely AV, et al. Reduced sleep spindles and spindle coherence in schizophrenia: mechanisms of impaired memory consolidation? Biol Psychiatry. 2012;71: 154–161. doi:10.1016/j.biopsych.2011.08.008

57. Cox R, Schapiro AC, Manoach DS, Stickgold R. Individual differences in frequency and topography of slow and fast sleep spindles. Front Hum Neurosci. 2017;11: 433. doi:10.3389/fnhum.2017.00433

58. Staresina BP, Bergmann TO, Bonnefond M, van der Meij R, Jensen O, Deuker L, et al. Hierarchical nesting of slow oscillations, spindles and ripples in the human hippocampus during sleep. Nature Neuroscience. 2015;18: 1679–1686. doi:10.1038/nn.4119

59. Denis D, Bottary R, Cunningham TJ, Davidson P, Yuksel C, Milad MR, et al. Slow Oscillation–Sleep Spindle Coupling Is Associated With Expectancy Measures of Fear Extinction Retention in Trauma-Exposed Individuals. Biological Psychiatry: Cognitive Neuroscience and Neuroimaging. 2025 [cited 10 Nov 2025]. doi:10.1016/j.bpsc.2025.08.009

60. Solano A, Riquelme LA, Perez-Chada D, Della-Maggiore V. Motor Learning Promotes the Coupling between Fast Spindles and Slow Oscillations Locally over the Contralateral Motor Network. Cerebral Cortex. 2022;32: 2493–2507. doi:10.1093/cercor/bhab360

61. Maris E, Oostenveld R. Nonparametric statistical testing of EEG- and MEG-data. J Neurosci Methods. 2007;164: 177–190. doi:10.1016/j.jneumeth.2007.03.024

62. Benjamini Y, Hochberg Y. Controlling the False Discovery Rate: A Practical and Powerful Approach to Multiple Testing. Journal of the Royal Statistical Society: Series B (Methodological). 1995;57: 289–300. doi:10.1111/j.2517-6161.1995.tb02031.x

## References

1. Denis D, Mylonas D, Poskanzer C, Bursal V, Payne JD, Stickgold R. Sleep Spindles Preferentially Consolidate Weakly Encoded Memories. J Neurosci. 2021;41: 4088–4099. doi:10.1523/JNEUROSCI.0818-20.2021

2. Gilson M, Tauste Campo A, Chen X, Thiele A, Deco, G. Nonparametric test for connectivity detection in multivariate autoregressive networks and application to multiunit activity data. Netw Neurosci. 2017;1:357-380. doi: 10.1162/netn_a_00019

